# A thermodynamical model of non-deterministic computation in cortical neural networks

**DOI:** 10.1101/2022.12.03.518978

**Authors:** Elizabeth A Stoll

**Author notes:** https://westerninstitute.org.

## Abstract

Neuronal populations in the cerebral cortex engage in probabilistic coding, effectively encoding the state of the surrounding environment with high accuracy and extraordinary energy efficiency. A new approach models the inherently probabilistic nature of cortical neuron signaling outcomes as a thermodynamic process of non-deterministic computation. A mean field approach is used, with the trial Hamiltonian maximizing free energy and minimizing the net quantity of temperature-entropy, compared with a reference Hamiltonian. Thermodynamic quantities are always conserved during the computation; free energy must be expended to produce information, and free energy is released during information compression, as correlations are identified between the encoding system and its surrounding environment. Due to the relationship between the Gibbs free energy equation and the Nernst equation, any increase in free energy is paired with a local decrease in membrane potential. As a result, this process of thermodynamic computation adjusts the likelihood of each neuron firing an action potential. This model shows that non-deterministic signaling outcomes can be achieved by noisy cortical neurons, through an energy-efficient computational process that involves optimally redistributing a Hamiltonian over some time evolution. Calculations demonstrate that the energy efficiency of the human brain is consistent with this model of non-deterministic computation, with net entropy production far too low to retain the assumptions of a classical system.

## I. INTRODUCTION

To compute the most likely state of the surrounding environment, a cortical neural network must select an optimal system state in the present context from a large probability distribution. Researchers have previously modeled this inherently probabilistic computation with Bayesian statistics [1], random-connection models [2], or fanofactor analysis of spike variance over time [3]. For individual neurons, the Hodgkin-Huxley equations provide a good approximation of firing patterns under steadystate conditions [4]. But channel leak and spontaneous subthreshold fluctuations in membrane potential significantly contribute to the likelihood of a given cell reaching action potential threshold [5-8]. Indeed, the relationship between membrane voltage, ion conductances, and channel activation, given by the Hodgkin-Huxley equations, is a classical limit that emerges from intrinsically stochastic processes [9-11]. Notably, cortical neurons actively maintain a coordinated ‘up-state’, allowing electrical noise to gate signaling outcomes [12]. Yet despite extensive literature on the statistical randomness of neuronal population coding and inter-spike variability, the mechanistic basis for achieving inherently probabilistic signaling outcomes across a cortical neural network is not well-understood.

Mean field theory has been usefully employed to model probabilistic coding in cortical neural networks [13, 14]. This methodology allows an exploration of the solution space, leading to the selection of a system state from a probability distribution [15]. At the mean field limit, the network achieves a fixed state, where excitatory and inhibitory contributions are balanced, so that fluctuations dominate the network level dynamics [16, 17]. As a result, the application of mean field theory has led to a better understanding of the contribution of internal membrane fluctuations to signaling outcomes [18] and how stochastic events shape network-level dynamics [19, 20].

Biological systems gradually achieve a more ordered configuration over time by identifying a more compatible state with their environment, simply by reducing predictive errors [21, 22]. This view, known as the free energy principle, asserts that learning systems strive toward ‘the minimization of surprise’, with new information continually prompting the revision of erroneous priors [23, 24]. Similarly, the concept of ‘free energy’ has been regularly employed in the machine learning field to solve optimization problems through a process of gradient descent [25, 26]. It should be noted however that ‘free energy’ is a statistical quantity, not a thermodynamic quantity, in these contexts. For the past forty years, researchers have striven to explain computation, in both biological and artificial neural networks, in terms of ‘selecting an optimal system state from a large probability distribution’ [27, 28]. Yet a mechanistic connection between noisy coding and energy efficiency has remained elusive.

This report presents a thermodynamic basis for meanfield theory, with the Hamiltonian being modeled not only as a computational quantity but also as an energetic quantity. Modeling cortical information processing as an iterative process of minimizing entropy and maximizing free energy ties together the bio-energetic efficiency of the system with the computational accuracy of the system. In this model, noise drives a non-deterministic computation, with the compression of information entropy paired with a release of free energy, which directly affects signaling outcomes. The extraordinary energy efficiency of the human brain is shown to be compatible with this model.

## II. METHODS

### A. A thermodynamic mean field model

A mechanistic process of thermodynamic computation is modeled with a Hamiltonian ⟨ *H*_*t*_⟩, which is the sum of all potential and kinetic energies in a non-equilibrium system. The Hamiltonian operates on a vector space, with some spectrum of eigenvalues, or possible outcomes, that can be obtained from a measurement. That measurement provides an exact solution for the Hamiltonian. This computational process resolves the amount of free energy available to the system *F*_*t*_, which is the total amount of energy in the system ⟨ *H*_*t*_⟨ less the temperature-entropy generated by the system *TS*_*t*_:

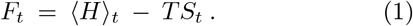

If no time has passed, or no interactions take place, the reference Hamiltonian *H*_0_ is the sum of all degrees of freedom *ξ*_*i*_ for all probabilistic components of the system:

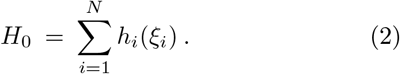

If some time has passed, or interactions have taken place, the Hamiltonian *H*_*t*_ of the system can be modeled as the mixed sum of all pairwise interactions:

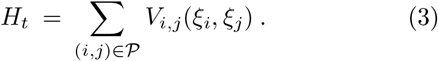

The mean field is then given by:

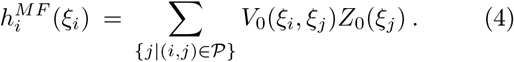

Where *V*_0_ represents the trace over *V*_*i,j*_ and *Z*_0_ represents the trace over 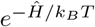. As the encoding System is perturbed, by interacting with its surrounding environment, the Hamiltonian evolves over time:

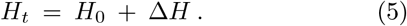

The Hamiltonian is dependent on an enormous number of contributing parameters. For this reason, it is computationally challenging to identify an exact solution, and variational methods in statistical physics use approximations to do so. Of course, different ‘measurements’ may yield different solutions, with different spectrums of eigenvalues. To model this variational outcome, we can employ a trial Hamiltonian:

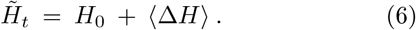

The original Hamiltonian has the same spectrum of eigenvalues as the trial Hamiltonian. However, the original Hamiltonian differs from the trial Hamiltonian by some positive value, such that:

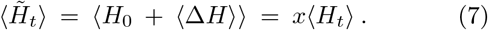

Since:

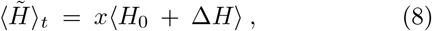

the free energy of the trial Hamiltonian must be greater than or equal to the free energy of the original Hamiltonian. This is known as the Bogoliubov inequality:

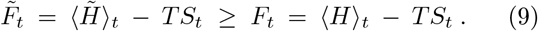

The computation results in energy being effectively redistributed across the system, with some trial Hamiltonian maximizing free energy availability. That trial Hamiltonian will be thermodynamically favored. The full account must always be balanced with the total amount of energy in the system, represented by the Hamiltonian, being the sum of all free energy and temperature-entropy.

### B. The entropy of the system

For a classical thermodynamic system, the macrostate of the system is a distribution of microstates, given by the Gibbs entropy formula. Here, *k*_*B*_ is the Boltzmann constant, *E*_*i*_ is the energy of microstate *i, p*_*i*_ is the probability that microstate occurs, and the Gibbs entropy of the system is given by *S*:

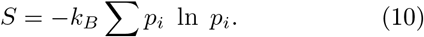

Yet the behavior of cortical neural networks can be better described as a statistical ensemble of microstates [1, 2]. And so, in this model, the macrostate of the system is formally described as a statistical ensemble of all component pure microstates, given by the von Neumann entropy formula:

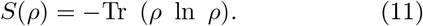

Here, entropy is a high-dimensional volume of possible system states, represented by the trace across a density matrix *ρ. ρ* is the sum of all mutually-orthogonal pure states *ρ*_*x*_, each occurring with some probability *p*_*x*_:

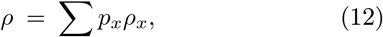

Each component state is described by a state vector *ψ*_*x*_. For example, the state of each neuron |*ψ*_*x*_*⟩*at time *t* is uncertain, described as having some probability of switching to an on-state (1) and some probability of remaining in an off-state (0). *ρ*_*x*_ is defined as the outer product of this finite dimensional vector space. The mixed sum of all these component pure states is therefore:

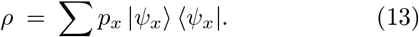

The cortical neural network is an open non-equilibrium thermodynamic system (System A), comprised of *N* units, each described by a state vector *ψ*_*Ax*_, operating within a surrounding environment (System B), comprised of *M* units, each with a state vector *ψ*_*Bx*_. Each system is described by a density matrix, or a mixed sum of orthonormal pure states. The system and its surrounding environment are initially uncorrelated with each other, and the combined system is created by the tensor product of the two density matrices:

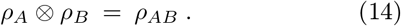

Each orthonormal pure state generates a Hilbert space, and any pure states that are identical cannot physically co-exist. Identifying non-distinguishable states (or linear correlations between pure states) will therefore compress the von Neumann entropy of the combined system. Any redundancies are eliminated during a linear transformation. And so, as the encoding Thermodynamic System ‘A’ is perturbed, by interacting with its surrounding environment, Thermodynamic System ‘B’, the density matrix undergoes a time evolution, from 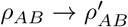:

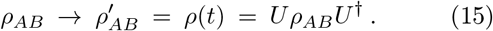

The unitary change in basis is provided by the time shift operator *U*:

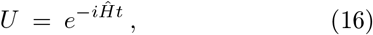

During this time evolution, correlations are identified between the two particle systems [29, 30]. If the systems are uncorrelated, the states are additive and the total entropy of the combined system remains unchanged. But if any component pure states between the two systems are correlated, entropy will be compressed. Here, entropy is additive in uncorrelated systems, but it is subtractive in correlated systems, as mutual redundancies in system states are recognized and reduced. This process of compression leads to the subadditivity rule:

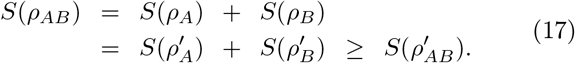

Thermodynamic System ‘A’ essentially solves a computationally complex problem by identifying correlations with its surrounding environment, Thermodynamic System ‘B’. An optimal system state in the present context is selected from a broad probability distribution, as information is compressed. The most thermodynamically favored and ‘optimal’ system state is the one that is both most correlated with the surrounding environment and most compatible with existing anatomical and physiological constraints.

### C. The free energy of the system

The Helmholtz equation can be used to calculate the net change in free energy over some period of time *t*. In a thermodynamic system that traps heat to accomplish work, the net change in Helmholtz free energy *F*_*t*_ is equivalent to the enthalpy *E*_*t*_, less the amount of temperatureentropy *TS*_*t*_ generated over that period of time:

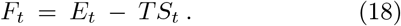

The Helmholtz free energy equation applies in contexts where pressure is not constant, but temperature is, while the Gibbs free energy equation applies in contexts where temperature is not constant, but pressure is. If the overall temperature and pressure of the system remain constant, the change in Helmholtz free energy *F*_*t*_ is equivalent to the change in Gibbs free energy *G*_*t*_:

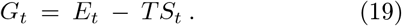

The Gibbs free energy of a given neuron is related to its membrane potential:

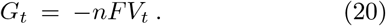

### D. The neuronal membrane potential

The Nernst equation calculates the temperature-dependent voltage shift in an electrochemical cell at thermodynamic equilibrium, based on the type and quantity of charge moving across the cellular membrane:

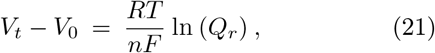

Where:

*V*_*t*_ is the electrochemical potential of the cell after some time *t* has passed (in volts),

*V*_0_ is the starting potential of the cell (in volts),

*R* is the universal gas constant (*R* = 8.314472 J/Kmol), *T* is the temperature in degrees Kelvin (*T* = 310.15 K under standard conditions),

*F* is the Faraday constant (*F* = 9.64853×10^4^ C/mol),

*n* is the number of electrons that are transferred during the reaction, and

*Q*_*r*_ is the reaction quotient, which defines the equilibrium potential of the reaction.

It is important to note the Nernst equation is specifically used for describing the resting potential of a neuron, not the action potential itself, which is a non-equilibrium process. This equation calculates the likelihood of a neuron firing an action potential upon some perturbance to the resting state (e.g. any event that increases the membrane potential or prompts inward sodium currents). Once the system has shifted away from equilibrium, the action potential goes to completion.

Thermodynamic systems in equilibrium are generally robust to shifting away from equilibrium, and the Nernst equation continues to describe the equilibrium state of the cell as that balance is restored, often returning the cell to a resting state. The Nernst equation only breaks down when the cell exits equilibrium – for example, when the sodium ion concentration inside the neuron increases and the equilibrium potential suddenly soars. That nonequilibrium state is the action potential, and it cannot be described by the Nernst equation.

Once a reaction proceeds, the Nernst equation cannot describe that non-equilibrium process. However, this computational process only affects the resting potential, so these time-dependent perturbations, occurring in the context of thermodynamic equilibrium, can be described by the Nernst equation.

## III. RESULTS

### A. A thermodynamic computation maximizes free energy and drives signaling outcomes

The total entropy generated by encoding System A, prior to compression, is defined as:

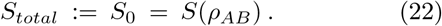

The net entropy generated by encoding System A, after compression, is defined as:

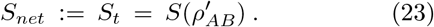

The change in entropy during compression is equivalent to the quantity of correlations identified between System A and System B:

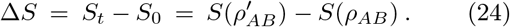

The quantity of entropy is *maximized* when the system state is completely random and all non-zero eigenvalues have equal probability *p*_*x*_. The randomness of information is *minimized* if a more predictable value is identified. In general, the von Neumann entropy of the system is less than maximal when some system states are more likely than others. *S*(*ρ*′_*AB*_) and *S*(*ρ*_*AB*_) reach equality only if no correlations are identified at all. During the time evolution, entropy is reduced, such that:

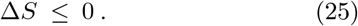

The energetic account must always be balanced, with the net amount of energy acquired by the system distributed toward either free energy or entropy:

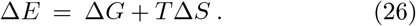

In order to balance the account, any loss of entropy during compression must be paired with a release of free energy, with Δ*G* = *G*_*t*_ − *G*_0_:

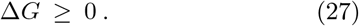

With these two values being equivalent:

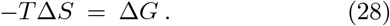

The compression of information entropy is paired with free energy release. This conservation law is known as the Landauer Principle [31-34]. This free energy is available to do *work*, allowing the system to attain a more ordered state, as it becomes less random and more compatible with its surrounding environment. This *work* involves building the membrane potential. The increase in Gibbs free energy, upon any reduction of uncertainty, is paired with an decrease in the neuronal membrane potential:

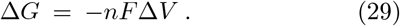

Information compression locally increases free energy and locally decreases the membrane potential. Neurons which have gained *certainty* during the computation will therefore gain free energy and move further away from action potential threshold, restoring their resting potential. By contrast, neurons which have gained *uncertainty* during the computation will lose free energy and move toward action potential threshold, increasing the likelihood of firing a signal.

If the overall temperature of the system remains the same, then information compression must both *decrease the distribution of possible system states* and *encode the most likely system state in a pattern of neural activity*. As a result, any reduction of uncertainty regarding the state of the surrounding environment will correspond to sparse but synchronous firing in neurons across the network.

In summary, the redistribution of energy during a thermodynamic computation, as the Hamiltonian is resolved, allows the heat-trapping system to physically instantiate the solution to a computational problem.

### B. Estimating net entropy production in the human brain

If the system gains internal heat over time *t*, then this energy can be distributed toward increasing entropy or increasing the amount of free energy available to do work. Minimizing the amount of entropy therefore leaves more free energy available to do work within the system. Since energy is always conserved, the account must always be balanced. Any reduction in entropy during the computation must be balanced by an increase in the amount of free energy that is available to do work. Conversely, any net increase in entropy must reduce the amount of free energy that is available to do work. If the brain does engage in non-deterministic computation, with a trial Hamiltonian maximizing free energy availability, then the brain should exhibit better-than-classical energy efficiency. The inefficiency of the system, or the net entropy production, can be calculated using empirical measures. The amount of temperature-entropy *T* Δ*S* produced over time *t* = Δ is equal to the net change in enthalpy Δ*E* less the change in available free energy Δ*G*:

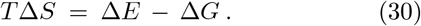

The enthalpy *E* is the total heat content of the system. If work is done *on* a system, Δ*E* is equal to the net change in internal energy of the system Δ*U plus* the net work done on the system Δ*W*. If work is done *by* the system, Δ*E* is given by the net change in internal energy of the system Δ*U minus* the net work done by the system Δ*W*. In the case of reversible changes to the quantities of either volume *V* or pressure *p* (nonelectrical work), Δ*W* = −*V* Δ*p*. For a cortical neural network – a far-from-equilibrium thermodynamical system of electrochemical cells, which actively traps energy to accomplish work – the quantity of Δ*E* is given by:

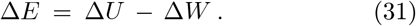

The change in the total heat content of the system Δ*E* is given by the net energy gained by the system Δ*U* subtracted by the amount of work completed by the system Δ*W* over some period of time *t*. The net energy Δ*U* is given by the quantity of thermal energy entering the system Δ*U*_*in*_ subtracted by the quantity of thermal energy exiting the system Δ*U*_*out*_ over time *t*. Combining Eqs. 30 and 31 yields:

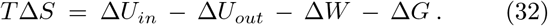

The net inefficiency of the system, given the temperature-entropy *T* Δ*S*, is equal to the net change in internal energy of the system Δ*U*, less the work done by the system Δ*W*, less the change in free energy Δ*G*.

Now that all energy in the system is accounted for, we can calculate values for Δ*U*_*in*_, Δ*U*_*out*_, Δ*W*, Δ*G*, and *T*, to estimate the quantity of Δ*S* generated by the human brain. The net change in internal energy of the system Δ*U* is equivalent to the caloric value supplied by the bloodstream to serve the neural network (provided by the energy input Δ*U*_*in*_) minus the amount of excess heat produced during that period of time (provided by the energy output Δ*U*_*out*_). To make a calculation based on neuronal signaling activity only, the change in free energy Δ*G* can be estimated as the quantity of energy released during the action potential, and the amount of work Δ*W* can be estimated as the quantity of ATP expended on setting up the electrochemical resting potential. The quantity of entropy Δ*S* produced in the human brain over time *t* can then be estimated by accounting for the constant overall temperature of the system, *T*:

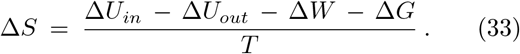

The energy consumed by the human brain over the course of a day is approximately 20% of oxygen intake and 20% of calories consumed by the body, a value that remains relatively constant regardless of variation in mental tasks or amount of motor activity [35, 36]. In adult males, this energetic usage is approximately 400 kilocalories or 1673.6 kJ per day. This estimated rate of Δ*U*_*in*_ is equal to 19.37 J/s or 19.37 Watts.

At rest, the human brain has an estimated metabolic rate of 3.5mL O_2_/100g/minute, with venous blood flow removing heat [37]. This rate yields a sustained jugular venous-to-arterial temperature difference of 0.3C [38, 39]. This value corresponds to an estimated heat production of approximately 6 J/kg/min [40, 41]. The rate of Δ*U*_*out*_ is therefore estimated to be 0.14 J/s or 0.14 Watts.

The amount of energy expended on work Δ*W* can be estimated by quantifying ATP turnover in the human brain. The quantity of ATP used on signaling processes in rat neocortex has been estimated at 21 *μ*mol g^−1^ min^−1^ [42], with experimental measurements of total ATP use approximating 30-50 *μ*mol g^−1^ min^−1^ [43-46], although estimates vary in both directions [47, 48]. Limiting the estimate to signaling processes only, the ATP turnover in neocortical grey matter is 0.35 *μ*mol g^−1^ s^−1^.

Assuming the human brain has a similar rate of ATP turnover to other mammals, the quantity of ATP used on signaling processes in the human brain can be calculated by estimating the total amount of neocortical grey matter. The size of the human brain by volume is rather variable, with a measured range of 1053-1499 cm^3^ in adult men and 975-1398 cm^3^ in adult women [49, 50]. The quantity of grey matter is 49.4% - 58.5% in adult men and 52.1% - 59.6% in adult women, both averaging 55% [51]. Since the average adult male brain is 1.4 kg [52], the approximate quantity of grey matter is 770 g, and so the estimated ATP turnover rate in this energetically expensive tissue is 0.27 mmol/s. In living cells, the hydrolysis of one ATP molecule releases 57 kJ/mol of energy. Given these values, the grey matter of the average adult male brain expends ATP on neuronal signaling processes at a rate of 15.36 J/s or 15.36 Watts. This is the estimated value of Δ*W*.

Here, Δ*U*_*in*_ is the amount of incoming caloric energy (19.37 J/s), Δ*U*_*out*_ is the heat loss from the system (0.14 J/s), Δ*W* is the amount of energy used to set up the electrochemical resting potential (15.36 J/s), and Δ*G* is the amount of free energy stored in the neuronal membrane that is released during the action potential (which we can assume to be negligible). Each parameter is provided as a rate of energy turnover in Joules per second, or Watts. Eq. 33 can be used to calculate the quantity of energy lost to entropy in the human brain every second, given by Δ*S*. Using the values for each parameter as described above, and a system temperature *T* equal to 37C or 310K, the value of Δ*S* over the course of *t=1s* can be calculated by substituting values into Eq. 33:

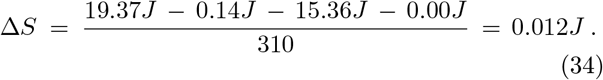

This exercise suggests the human brain is 99.9% energy-efficient, an estimate that corresponds well with other models demonstrating the remarkable energetic efficiency of cortical neurons within the central nervous system [42-48]. The high rate of energetic turnover in these cells should contribute to significant quantities of entropy and heat production, yet this seems not to be the case. There is far too much ordered work happening here. In addition to electrochemical signaling, neurons undergo anabolic metabolism, gene transcription, protein manufacture, post-translational modification, protein transport, and membrane remodeling, all processes which require energetic expenditure. Notably, these activity-dependent tasks maintain the ordered state of the neuron and ensure the physical structure retains an optimal configuration to encode information within the neural network. These non-signaling tasks are not included in the budget for ATP turnover given here, and may be reasonably expected to further increase the energy expenditure of cortical neurons. In addition, the 45% of brain weight which is not cortical grey matter is not included in this estimate, contributing to 20% of the total ATP turnover of the brain, or an additional 3.8 J/s. As a result, the total amount of energy expended on performing work appears to exactly match total caloric uptake – leading to the astonishing conclusion that the human brain is nearly 100% energy efficient.

And so, with excellent estimates for caloric intake, heat output, ATP turnover, and temperature – based on empirical data – the quantity of *energy* expended to complete *work* within the mammalian central nervous system is found to be curiously efficient. As a result of this extraordinary efficiency, the estimated quantity of *entropy* produced by the human brain is far too low to retain the assumptions of a classical system.

It seems highly unlikely that a physical system whose primary job is to process information creates no physical information or entropy at all. Instead, this model shows how a heat-trapping system might cyclically generate and compress information entropy, recovering free energy as that thermodynamic quantity of information is parsed for *correlations, consistencies*, or *predictive value*.

### C. Predictions

The theoretical model presented here results in an energy-efficient, non-deterministic, system-wide computation. As correlations are identified, and information entropy is physically compressed, the thermodynamic computing system takes on a more ordered state and becomes more compatible with its surrounding environment. This theoretical framework therefore offers a putative mechanistic link between non-deterministic computation and extraordinary energy efficiency in the mammalian brain.

This new theoretical framework for modeling non-deterministic computation in cortical neural networks makes some specific predictions with regard to the wavelength of thermal free energy released upon information compression [53], and the contribution of these localized thermal fluctuations to cortical neuron signaling outcomes [54]. This approach also makes specific predictions about the expected effects of electromagnetic stimulation and pharmacological interventions on perceptual content [55]. Some further predictions of the theory, prompted by the present model, include:

#### 1. Cortical neurons should meet the criteria for a quantum system

In this new theoretical model, cortical neurons redistribute a Hamiltonian operator to minimize entropy and maximize free energy, with this computational process driving signaling outcomes. This is explicitly a model of quantum computation, with probabilistic coding cyclically generating and compressing quantum information. Certainly doubt has been cast on the hypothesis that the brain is a quantum system [56]. If this hypothesis is true, then empirical measures of coulomb scattering profiles, decoherence timescales, ionization dynamics and dissipation rates should meet the Tegmark criteria for a quantum system: “If *τ* _dyn_ ≪ *τ* _dec_, we are dealing with a true quantum system, since its superpositions can persist long enough to be dynamically important. If *τ* _dyn_ ≫ *τ* _diss_, it is hardly meaningful to view it as an independent system at all, since its internal forces are so weak that they are dwarfed by the effects of the surroundings. In the intermediate case where *τ* _dec_ ≪ *τ* _dyn_ *< τ* _diss_, we have a familiar classical system.” Further studies should be conducted to ascertain the exact values for each of these parameters in cortical neurons.

#### 2. Cortical neurons should demonstrate exceptional energy efficiency

In this new theoretical model, a cortical neural network selects an optimal system state in the present context from a large probability distribution, in a process of non-deterministic computation. This naturally leads to the compression of information entropy and the release of free energy. If cortical neurons are indeed able to recover the energy that is normally dissipated irreversibly toward the production of entropy, their net entropy production should be well below classical expectations for the amount of work being completed. If cortical neurons do undergo a physical process of non-deterministic computation, then empirical studies should confirm these cells to be significantly more energy-efficient than neurons in spinal reflex circuits, which have purely deterministic signaling outputs. This prediction can be tested by comparing ATP turnover in cells of similar size, with similar firing rates and similar levels of gene expression, protein turnover, and intracellular transport. If cortical neurons are classical computational units, obeying the null hypothesis, then these cells should exhibit purely classical energy efficiency.

#### 3. Artificial neural networks with the physical properties of a cortical neural network should achieve spontaneous unprogrammed exploratory behavior

One challenge for both organisms and robotics is spontaneously exploring the local environment in search of predictive value, without being explicitly programmed to do so [57]. Recent advances in state-of-the-art robotics have involved introducing a library of robot action primitives, parameterized by arguments which are gradually adopted under a reinforcement learning policy [58]. In classical computing architectures, there simply cannot be spontaneous acceleration from rest without programmed priors and policies. By contrast, in this model of quantum computing architecture, energy is periodically redistributed around the system, to achieve directed work - including synaptic remodeling and driving the action of the body. For this reason, far-from-equilibrium systems that trap *heat* to do *work* can spontaneously explore their environment and gain knowledge about it through this method of non-deterministic computation. This theory predicts that such unprogrammed exploratory behavior can never be achieved in classical computing architecture, because the Hamiltonian cannot be redistributed; however, robotic hardware with similar anatomical and physiological properties should be able to achieve spontaneous learning by engaging with their local environment.

#### 4. Artificial neural networks with the physical properties of a cortical neural network should require fewer time and energy resources to solve computationally complex problems

Another challenge for both organisms and robotics is minimizing the time and energy resources utilized while solving decision problems. The system must encode the most likely state of the surrounding environment, using one or more sensory modalities impinging on a layered neural network architecture, then make a decision on the appropriate response in that context. In classical computing architecture, computationally complex problems can be solved through brute force methods [59] or cascading classifiers [60]. Alternatively a large solution space can be explored, with a minimization of energy expenditure achieved through gradient descent [25, 26]. In this model of quantum computing architecture, time and energy are uncertain until these variables are resolved into a mutually compatible state for all computational units within the system. For this reason, this theory predicts that fewer time and energy resources should be needed to solve computationally complex problems using this method of thermodynamic computation. However, robotic hardware with similar anatomical and physiological properties should exhibit biologically-comparable time and energy efficiency during decision problems, if the Hamiltonian is being effectively redistributed.

## IV. DISCUSSION

The extraordinary energetic efficiency of the central nervous system has been noted, particularly among theorists who query whether this competence is intrinsically linked to the production of information entropy [61, 62] or the exascale computing capacity of the brain [63, 64]. In this report, energy is indeed being expended on the production of information – but rather than being an abstract quantity, or a quantity that is irreversibly lost, this quantity is released back into the system as free energy as information is physically compressed, in accordance with the Landauer principle.

This approach models how cortical neural networks produce entropy in a thermodynamic sense, then evaluates whether information processing emerges from such a lean assumption. Indeed, all thermodynamic systems must create entropy, with that quantity related to the pressure, temperature, density, and volume of the system. If a hot, dense, non-heat-dissipating system appears to be nearly 100% energy-efficient, the sensible logic is *not* that entropy is not being produced at all, but rather that correlations are being extracted and information is being compressed, in accordance with physical laws. This report demonstrates that information generation and compression can occur in far-from-equilibrium systems, with the system naturally encoding the likely state of its surrounding environment, as it takes on a more ordered state over time.

This model of neural computation both vindicates and elaborates Friston’s free energy principle [21-24], providing a thermodynamic basis for the reduction of ‘surprise’ during predictive processing. While prior efforts used statistical methods to model the inherently probabilistic patterns of cortical neural network activity [1-3, 18], the present model usefully shows how energy-efficient non-deterministic computation might be achieved by following thermodynamic laws.

Here, employing the laws of information thermodynamics yields a mechanistic connection between probabilistic coding and the free energy principle in cortical neural networks. Moving forward, exploring the relationship between energetic efficiency and non-deterministic computation may prove useful to both the field of neuroscience and the field of computational physics.

## ACKNOWLEDGMENTS

The author received support for this work from the Western Institute for Advanced Study, with generous donations from Jason Palmer, Bala Parthasarathy, and Vanguard Charitable.

